# Long-read sequencing identified a causal structural variant in an exome-negative case and enabled preimplantation genetic diagnosis

**DOI:** 10.1101/326496

**Authors:** Hefan Miao, Jiapeng Zhou, Qi Yang, Fan Liang, Depeng Wang, Na Ma, Bodi Gao, Juan Du, Ge Lin, Kai Wang, Qianjun Zhang

## Abstract

For a proportion of individuals judged clinically to have a recessive Mendelian disease, only one pathogenic variant can be found from clinical whole exome sequencing (WES), posing a challenge to genetic diagnosis and genetic counseling. Here we describe a case study, where WES identified only one pathogenic variant for an individual suspected to have glycogen storage disease type Ia (GSD-Ia), which is an autosomal recessive disease caused by bi-allelic mutations in the *G6PC* gene. Through Nanopore long-read whole-genome sequencing, we identified a 7kb deletion covering two exons on the other allele, suggesting that complex structural variants (SVs) may explain a fraction of cases when the second pathogenic allele is missing from WES on recessive diseases. Both breakpoints of the deletion are within Alu elements, and we designed Sanger sequencing and quantitative PCR assays based on the breakpoints for preimplantation genetic diagnosis (PGD) for the family planning on another child. Four embryos were obtained after in vitro fertilization (IVF), and an embryo without deletion in *G6PC* was transplanted after PGD and was confirmed by prenatal diagnosis, postnatal diagnosis, and subsequent lack of disease symptoms after birth. In summary, we present one of the first examples of using long-read sequencing to identify causal yet complex SVs in exome-negative patients, which subsequently enabled successful personalized PGD.

## Abbreviations

ACMG: The American College of Medical Genetics and Genomics
GSD: Glycogen storage disease
GSD-Ia: Glycogen storage disease type Ia
G6PC: glucose-6-phosphatase
HGMD: Human gene mutation database
IVF: In-vitro fertilization
PCR: Polymerase chain reaction
PGD: Preimplantation genetic diagnosis
WES: Whole exome sequencing

## Introduction

Whole exome sequencing (WES) is now widely used in genetic testing on patients who are suspected or have been clinically demonstrated to have genetic disorders. However, a large proportion (~60-70%) of patients judged clinically to have a Mendelian disease receive negative results on WES with current Illumina short-read sequencing technology ^1–5^. Compared to WES, the use of whole-genome sequencing (WGS) does not appear to significantly improve the diagnostic yield ^6,7^ or present much economic advantage ^8^. Therefore, WES/WGS-negative cases pose a significant challenge to the clinical diagnosis of genetic diseases. Several reasons may explain the lack of positive findings, such as the inefficiency of template DNA capture, the biases in sequencing coverage, the failure to call causal variants from data, the inability to catalog all functional variants (especially non-coding variants), the incorrect clinical interpretation of genetic variants, the possibility of complicated oligogenic disease in certain patients and the possibility of disease causal mechanisms due to somatic or epigenetic origin. Among these reasons, the limited ability to interrogate repeat elements such as tandem repeats ^9^ and structural variants (SVs) ^10^ may play important roles. In fact, several published studies demonstrated examples where disease causal SVs were missed by short-read WES/WGS ^11^, or that certain classes of disease causal repeats failed to be identified by WES/WGS ^12^.

Long-read sequencing technologies, such as the 10X Genomics linked-read sequencing, the Oxford Nanopore Technologies (ONT) and PacBio single molecule real-time (SMRT) sequencing, offer complementary strengths to traditional WES/WGS based on short-read sequencing. Long-read sequencing can produce read length (typically 1,000 bp or longer) that is far higher than the 100-150bp produced by short-read sequencing, therefore allowing for the resolution of breakpoints of complex SVs ^13^ or the detection of long tandem repeats ^14,15^. In particular, recent *de novo* human genome assemblies via long-read sequencing have revealed tens of thousands of SVs per genome, several times more than previously observed via WGS, suggesting an underestimation of the extent and complexity of SVs ^16–19^.

In the present study, we applied a long-read whole-genome sequencing to yield the genetic diagnosis of glycogen storage disease type Ia (GSD-Ia) in a patient whose causal variants were unsolved by Sanger sequencing and WES. Glycogen storage disease type I (GSD-I) is a group of autosomal recessive metabolic diseases caused by defects in the glucose-6-phosphatase (G6Pase) complex, with an overall incidence of approximately 1:20,000-40,000 cases per live birth ^20,21^. The most common form, GSD-Ia, represents more than 80% of GSD-I cases ^22^. Mutations in the *G6PC* gene have been found to be the cause of this disease. The *G6PC* comprises of five exons on chromosome 17q21, and encodes a 35 kD monomeric protein, the G6Pase catalytic subunit, which plays a role in the endoplasmic reticulum ^23,24^. Through Sanger sequencing and WES, we were able to identify one deleterious mutation in the proband, yet we suspected the presence of a complex SV due to the observation of Mendelian inconsistency in the family. As a result of long-read sequencing, we made a positive diagnosis of GSD-Ia on the patient and accurately identified the breakpoints of a causal SV in the other allele of the G6PC gene, which further guided genetic counseling in the family and enabled a successful preimplantation genetic diagnosis (PGD) for in vitro fertilization (IVF) on the family.

## Results

### Clinical examination

We were presented with a 12 year-old boy with hepatosplenomegaly and growth retardation at the Xiangya Hospital of Central South University, Hunan, China in 2017 (Figure 1A). The clinical features include a rounded doll’s face, fatty cheeks and protuberant abdomen (Figure 1B). Based on examination on the proband’s skeletal development by X-rays, his hip and wrist showed osteoporosis (Figure 1C, 1D). In the upper liver, right intercostal midline sixth intercostal, liver rib length is 51 mm, thickness is 37 mm. Maximum oblique diameter of right liver is 159 mm, suggesting severe liver enlargement (Figure 1D). Spleen is swollen to a thickness of 34 mm.

**Figure 1.**
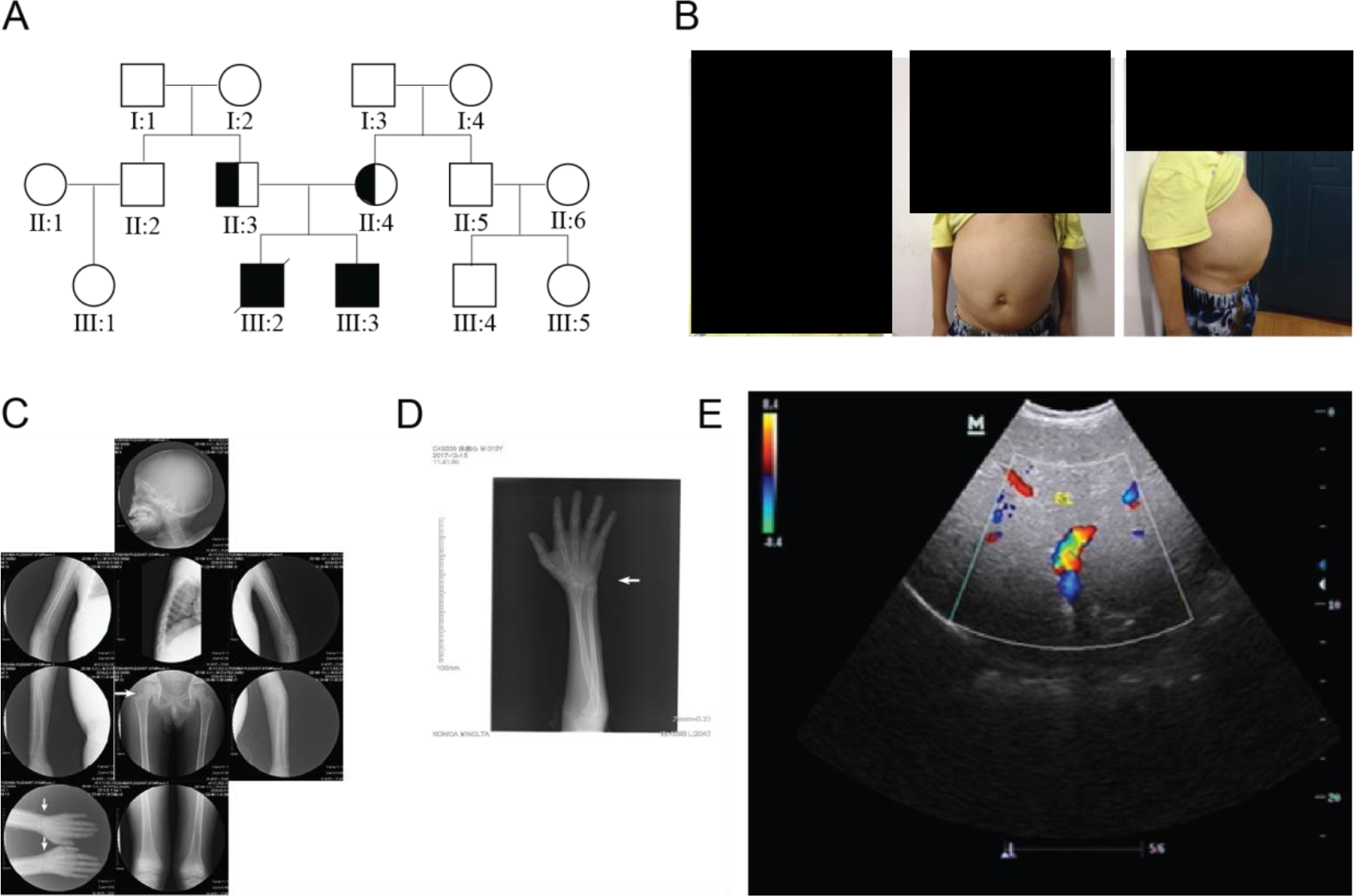
Clinical characteristics of the proband. (A) Pedigree of the family. III:3 represents the proband, whose older brother (III:2) has decreased. (B) The clinical features include a rounded doll’s face, fatty cheeks and protuberant abdomen. (C) X-ray films of the whole body of the patient. White arrows mark areas with obvious osteoporosis. (D) Focused view of X-ray film on the hand of the proband, where the wrist marked by white arrows has obvious osteoporosis. (E) Image of type-B ultrasonic on the proband shows severe liver enlargement. Blue color: The blood flow away from the detector of ultrasound B-mode scanner; Red color: The blood flow to the detector of ultrasound B-mode scanner.

Additional biochemical assays were performed on the proband (Table 1). His fasting blood glucose value was 0.17 mmol/l, which had reached a dangerously low value. Levels of cholesterol, triglycerides and chlorinum were abnormal. Investigation of liver function showed elevated aminotransferases (AST, ALT) and other biochemical abnormalities. Serum copper and ceruloplasmin were at normal levels. Altogether, these clinical and biochemical examinations indicated that the boy is likely to be affected with GSD-Ia.

**Table 1.** Biochemical indicators of the proband suggest a probable diagnosis of GSD-Ia.

| Parameter | Tested value | Reference range | Result |
| --- | --- | --- | --- |
| Glucose (mmol/L) | 0.17↓ | 3.6-6.1 | Abnormal |
| CO2 (mmol/L) | 7.1↓ | 19-33 | Abnormal |
| Sodium (mmol/L) | 132.7 | 135-153 |  |
| Chlorinum (mmol/L) | 89.2↓ | 96-108 | Abnormal |
| Calcium (mmol/L) | 2.65 | 2-2.6 |  |
| Total bile acid (μmol/L) | 16.7↑ | 0-12 | Abnormal |
| ALT (IU/L) | 257.4↑ | 7-56 | Abnormal |
| AST (IU/L) | 357.6↑ | 0-40 | Abnormal |
| Triglyceride (mmol/L) | 5.74↑ | 0.52-1.56 | Abnormal |
| HDL (mmol/L) | 2.01↑ | 0.88-1.76 | Abnormal |

Abbreviations: ALT, alanine aminotransferase; AST, aspartate aminotransferase; HDL, high-density lipoproteins.

**Table 2.**
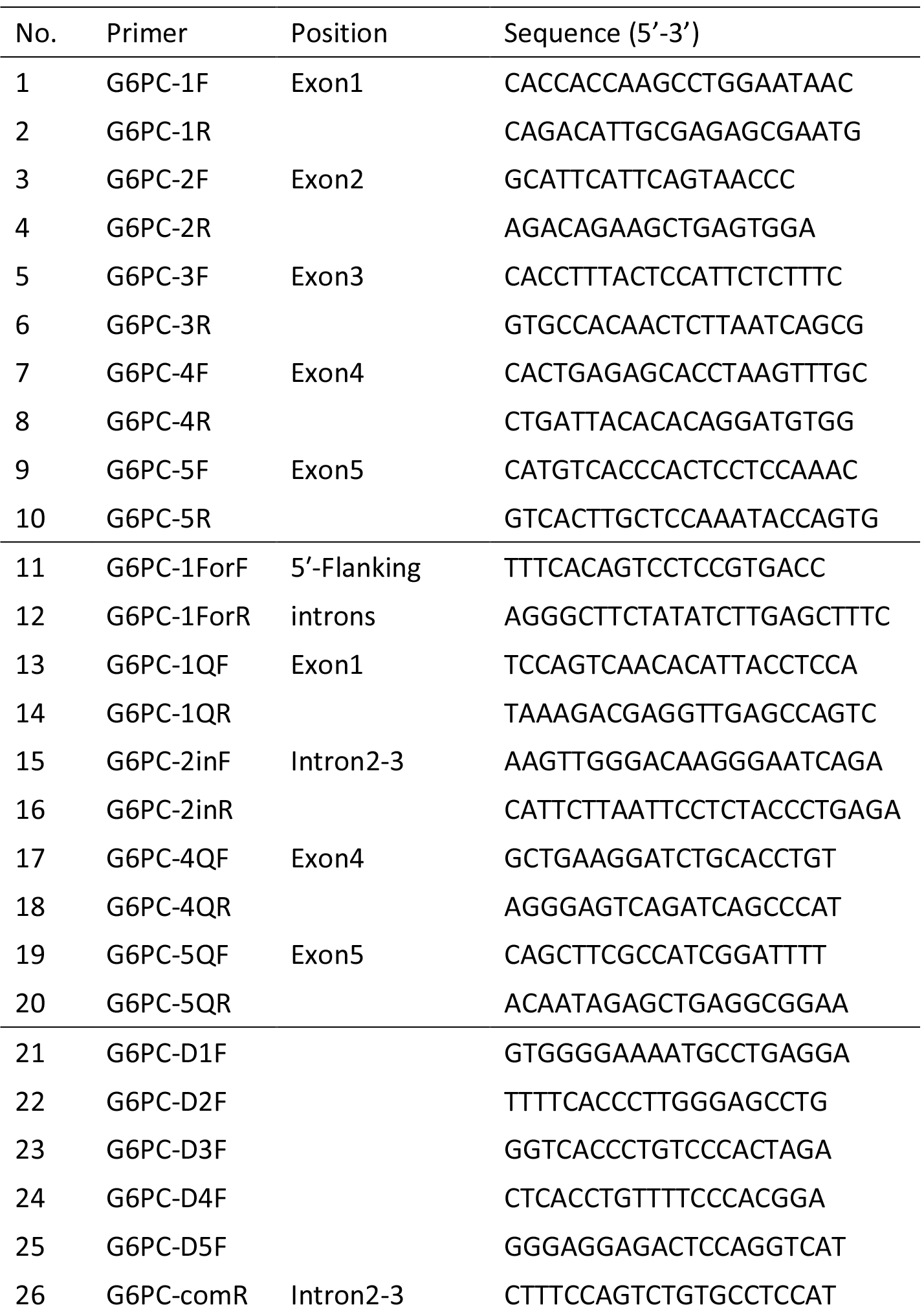
Primers for Sanger sequencing. No.1-10: Exon primers; No.11-20: Quantitative real-time primers; No.21-26: Breakpoint primers.

The parents were non-consanguineous, and neither has any symptom of GSD-Ia (Figure 1A). They came to Xiangya Hospital to seek help to obtain a genetic diagnosis and plan to have another child by in vitro fertilization (IVF). Given that GSD-Ia is a recessive disease, we hypothesized that the proband carries bi-allelic *G6PC* mutations inherited from the father and mother, respectively. Yielding a confirmed genetic diagnosis and determining the exact disease causal variants are necessary to perform preimplantation genetic diagnosis (PGD) from in vitro fertilization (IVF).

### Whole-exome and Sanger sequencing identified one pathogenic variant

To confirm the clinical diagnosis, we conducted clinical exome sequencing on the proband. A novel missense mutation of the G6PC gene (c.326G>A) was identified by WES (Figure 2A), which appears to be homozygous and affects a highly conserved position in the protein sequence (Figure 2B). Bioinformatics analysis by InterVar ^25^ and manual examination of the ACMG-AMP 2015 guidelines ^26^ determined the mutation to be likely pathogenic. We further validated the mutation by Sanger sequencing on all family members (Figure 2C). However, the mutation was not detected in the father and was present in a heterozygous state in the mother (Figure 2C). In order to examine whether germline mosaicism is present in the father, the DNA of the father’s sperm was sequenced, but the results were largely inconclusive as a small peak for A allele and an even smaller peak for C allele is present at the c.326 position (Figure S1).

**Figure 2.**
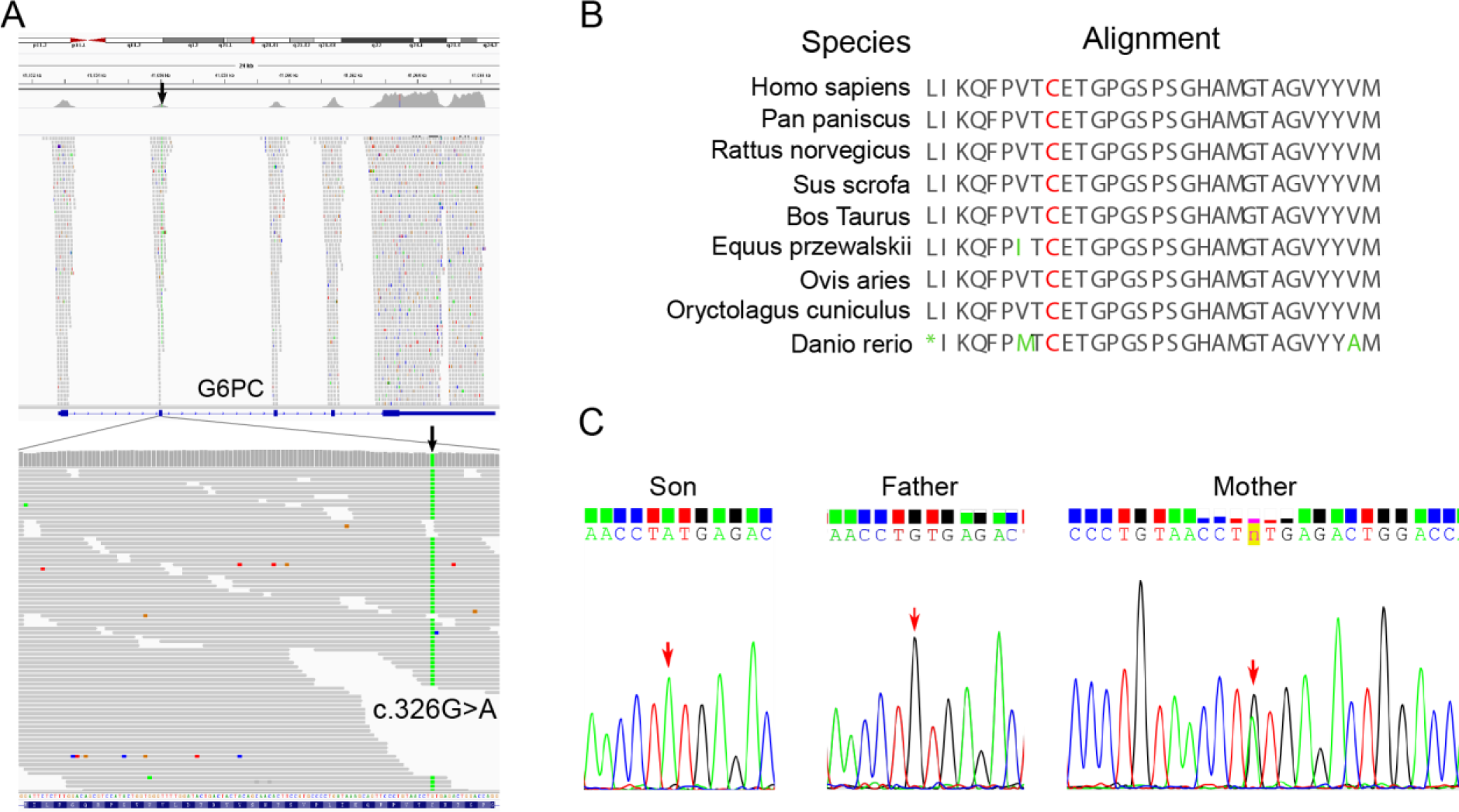
Identification of a c.326G>A missense mutation in the *G6PC* gene. (A) Whole-exome sequencing identified a homozygous c.326G>A missense variant in exon 2 of the *G6PC* gene. (B) The amino acid 109 (marked by red color) affected by c.326G>A is highly conserved across different species. (C) Sanger sequencing on the pedigree showed that the father does not carry the c.326G>A missense variant and that the mother carries a heterozygous c.326G>A missense variant.

### Long-read sequencing identified a structural variant in the other allele

To evaluate whether a structural variant is present in the proband that masks the c.326G>A mutation as homozygous, we carried out long-read whole-genome sequencing on the proband using the Oxford Nanopore technology. We generated 2,251,269 base-called reads containing 35,595,548,336 bases (~12X whole-genome coverage) with an average read length of 16,579 bp. Using the long-read sequencing data, a novel deletion on chr17 g.41049904_41057049del7146 (GRCh37) was detected in one allele of G6PC (Figure 3A), and the known heterozygous point mutation c.326G>A was detected in the other allele. This deletion was supported by four reads, though with slightly discordant breakpoints due to possible alignment errors. The deletion completely covers the first two exons of the G6PC gene, thus resulting in loss of function.

**Figure 3.**
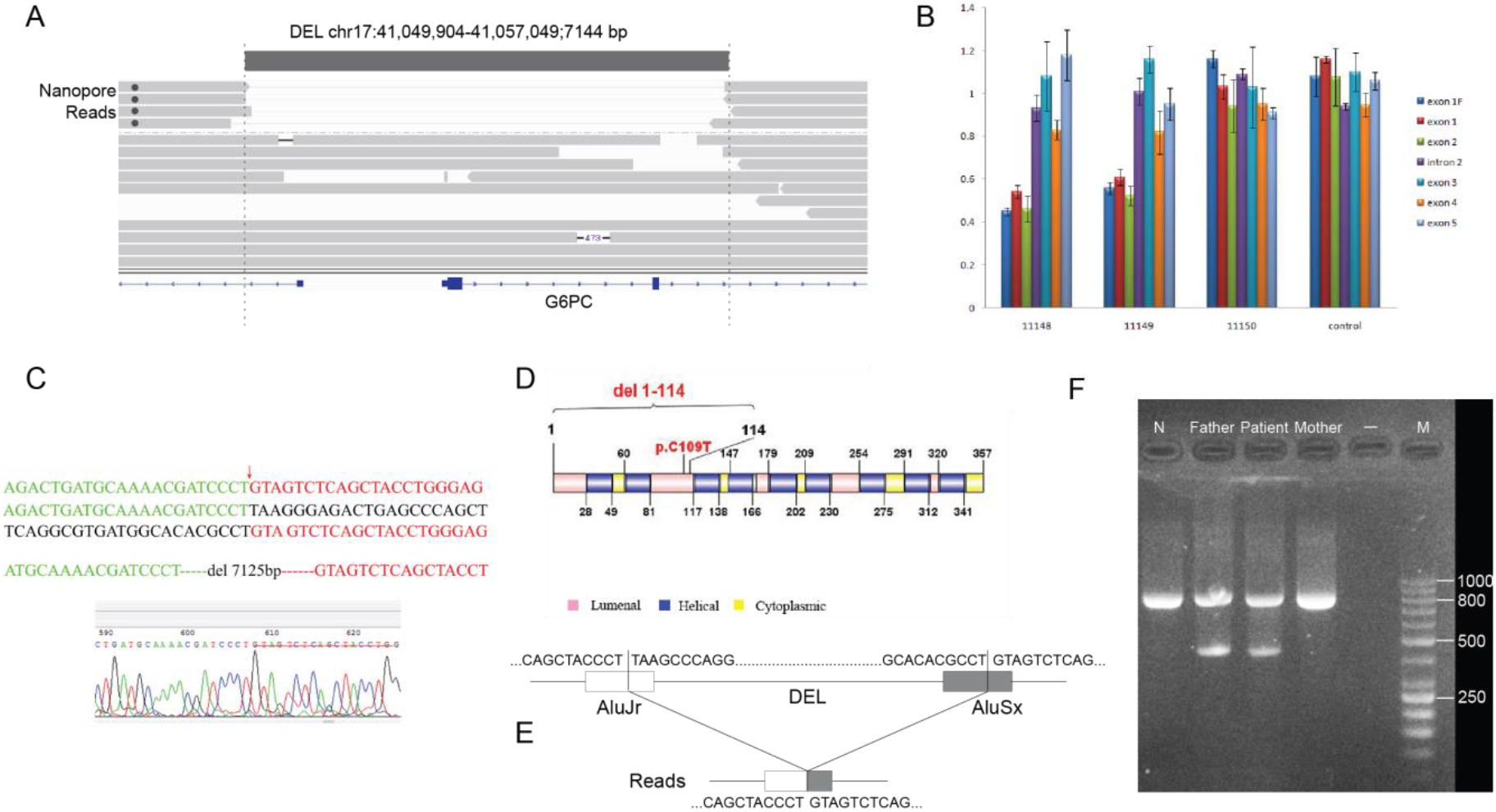
Long-read sequencing identified a deletion in the *G6PC* gene. (A) IGV screen shot of reads at the *G6PC* locus. Four reads carry a deletion (chr17 g.41049904_41057049del7125 that starts from the first intron of the *LINC00671* gene to intron 2 of the *G6PC* gene. (B) Quantitative PCR validation of the deletion in the trio. Relative quantitation (RQ) of copy number was analyzed by the ΔΔCT method, and error bars represent standard deviation. The deletion includes exon 1F (5′-Flanking introns), exon 1 and exon 2, and the patient and his father are mutation carriers while his mother is normal. (C) Sanger validation of the deletion breakpoints. The first sequence shows the mutated genomic segment, while the second and third sequences show expected genomic segments if deletion is not present. The red arrow refers to the breakpoint, and a 7125 bp sequence is deleted based on the human reference genome (GRCh37). (D) Depiction of the protein domains that were targeted by the non-synonymous mutation and the 7kb deletion. (E) Illustration of the genomic contexts of the two breakpoints, which are both located in known Alu elements. (F) Gel electrophoresis of the PCR product designed to detect the deletion. The lane marked with M represent GeneRuler 50 bp DNA Ladder (Thermo Scientific^™^), and all lanes (except “-” lane) include an ~800bp internal control (β-Globin gene). A 418bp fragment can be amplified from the father and the proband.

By quantitative RT-PCR, we estimated the copy number of each exon of the G6PC gene from the proband and his parents (Figure 3B). The copy numbers of deletion exon 1F (5′-Flanking region), exon 1 and exon 2 were only about half of the value of the control group, which indicated that the patient and his father both carry a heterozygous deletion.

To further clarify the location of the breakpoint of this large deletion, we designed six breakpoint primers (Table 1) to amplify and sequence the suspected breakpoint region. The result confirmed our prediction and found the breakpoints precisely at chr17:41049879 and chr17:41057003 (Figure 3C). These breakpoints were only 20-50bp different from the predictions from the long-read sequencing data. This large deletion (chr17 g.41049879_41057003del7125) was thus 7125 bp in length and contains the 5′ regulatory sequence as well as exon 1, intron 1, exon 2 and partial intron 2 of the G6PC gene (Figure 3D). Motif analysis was performed and found that the 5′ breakpoint was located in AluJr and the 3′ breakpoint in AluSx (Figure 3E). The Alu family members have a sequence similarity of over 87% and cover 11% of the human genome ^27^, and Alu-Alu recombination usually produces a fragment deletion via Alu recombination-mediated deletion (ARMD)^28,29^. Note that WES missed this deletion and judged the heterozygous missense mutation as homozygous, since we were unable to observe any obvious coverage differences between the several exons in the gene from WES (Figure 2A). Therefore, the patient inherited the missense mutation (c.326G>A) from his mother and inherited the deletion mutation (chr17 g.41049879_41057003del7125) from his father. We further designed a PCR-based assay to easily differentiate samples with and without deletion using a set of primers (G6PC-Del-F/G6PC-Del-R), which generates a 418bp target fragments in individuals carrying the deletion (Figure 3F).

### Preimplantation genetic diagnosis and implantation of the embryo

In order to help the family to plan for another child, a reproductive intervention was carried out by in vitro fertilization with preimplantation genetic diagnosis (PGD). To avoid the allelic drop-out (ADO), microsatellite markers D17S760 (location 5′ of G6PC, −0.9M), DS17S793 (location 5′ of G6PC, −0.7M) and DS17S951 (location 3′ of G6PC, 0.8M) in linkage with the breakpoint junction and the point mutations were tested. Four embryos developed to blastocysts, and were collected and biopsied. The deletion mutation (chr17 g.41049879_41057003del7125) was not detected in four embryos, and we ruled out the possibility of allelic drop-out via linkage analysis by microsatellite markers. However, all the four embryos were identified to be carriers of the missense mutation (c.326G>A). Data on the STR sites D17S760, DS17S793 and DS17S951 also indicated that the embryo inherited the maternal risk chromosome (Table S2). Furthermore, three alleles of D17S760 were found in embryo No. 2, suggesting that it may be partial trisomy in chromosome 17. Considering the state of the four embryos comprehensively, embryo No. 1 was implanted (Table S1).

After the implantation of the embryo, the patient’s mother succeeded in pregnancy and came to our hospital for a review at the 19+ week of pregnancy. We obtained amniotic fluid cells of the fetus to extract DNA. The genetic testing confirmed that the fetus is a carrier of the missense mutation (c.326G>A), and does not inherit the deleterious deletion. The newborn was revisited in our hospital in December 2017. The baby had a fasting blood glucose level of 5.5 mmol/L and her B-ultrasonogram showed that the liver and kidneys were normal (Figure S2), confirming that she was not affected with GSD-Ia.

## Discussion

In the current study, we performed genetic diagnosis on an affected subject with suspected GSD-Ia through clinical whole-exome sequencing and long-read whole-genome sequencing. The proband was misidentified as a homozygote for the c.326G>A mutation by WES, but later confirmed to be a compound heterozygous carrier of the c.326G>A mutation and a 7kb deletion spanning this point mutation. The missense mutation and the deletion were inherited from the mother and father, respectively. Therefore, through combined exome sequencing and long-read whole-genome sequencing, we yielded a definitive genetic diagnosis on the proband, and used this information to design assays to enable successful personalized preimplantation genetic diagnosis following IVF.

The human G6PC is a single-copy gene that contains five exons and spans 12.5 kb of DNA on chromosome 17q21 ^30^. The G6PC gene encodes G6Pase which is a 357 amino acid protein anchored to the endoplasmic reticulum (ER) membrane with nine transmembrane domains ^31^. The amino-terminus of the protein lies in the ER lumen with the enzymatic active site and the carboxyl-terminus in the cellular cytoplasm ^20,32^. Based on the predicted structure on UniProt (http://www.uniprot.org/uniprot/P35575), the novel point mutation (c.326G>A; p.C109Y) detected in this study is located in the lumen of the ER and the deletion fragment (chr17 g.41049879_41057003del7125) contains at least five transmembrane domains. Since the deletion spans the starting codon of the protein, the allele carrying the deletion should not to be transcribed or translated.

To date, differential diagnosis of GSD generally relies on the molecular analysis and has replaced the traditional liver biopsy ^22,32^. Detection and analysis of suspicious disease-causing mutations have become a powerful tool for differential diagnosis of GSD and can guide the implementation of PGD and the personalized treatment even further. WES has gradually become a powerful means by which clinicians and scientists can detect the underlying cause of various genetic diseases. However, a major shortcoming of WES is uneven coverage of sequence reads over the capture regions, contributing to many low coverage regions, which hinders accurate variant calling ^33^. This challenge did not affect our study per se, since we were able to identify the c.326G>A mutation from WES accurately, but the uneven coverage from exome sequencing prevented us from finding the deletion covering two exons. Nevertheless, given that this mutation is a very rare mutation (not documented in public databases) and that there is no known consanguinity in the family, it is unlikely that the proband inherits both alleles from parents. Initially we suspected that the father carries germline mosaicism, yet sperm sequencing on the father did not fully resolve the question (Figure S1). A low coverage long-read whole-genome sequencing resolved this issue. A large deletion (initially designated as chr17:g.41049904_41057049del7146 based on alignment) was detected in the proband and his father. Then we conducted the Sanger sequencing and the quantitative RT-PCR to validate this result and refine the exact position of the breakpoint junction (chr17:g.41049879_41057003del7125). The slight inconsistence of the deletion breakpoints between initial long-read sequencing and subsequent Sanger sequencing were likely due to the higher error rate of long-read sequencing and the imperfect alignment of the reads. Long-read sequencing can identify complex SVs effectively, thus compensated for the shortcomings of WES and avoided a misdiagnosis and potential failure of PGD.

In summary, we present one of the first examples of using long-read sequencing to identify causal yet complex structural variants (SVs) in exome-negative patients, which subsequently enabled successful personalized preimplantation genetic diagnosis. Our study suggests that long-read sequencing offers a means to discover overlooked genetic variation in patients undiagnosed or misdiagnosed by short-read sequencing, and may potentially improve diagnostic yields in clinical settings, especially when only one pathogenic mutation is found in an affected individual suspected to carry a recessive disease.

## Materials and Methods

### Patient characteristics

The study was approved by the CITIC-Xiangya Reproductive and Genetics Hospital, Central South University. The collection and use of tissues followed procedures that are in accordance with ethical standards as formulated in the Helsinki Declaration, and informed consent was obtained from the study participants. The proband was a 12 year old boy with hepatosplenomegaly and growth retardation, and was diagnosed with possible GSD-Ia in Xiangya Hospital of Central South University in 2017. The parents of the proband came to Xiangya Hospital to seek help to have another child via IVF, but genetic testing by Sanger sequencing and whole-exome sequencing (WES) failed to identify a definitive genetic cause of the disease in the family. Indeed, a homozygous mutation of c.326G>A (p.C109Y) in exon 2 of the G6PC gene was identified via WES and was interpreted to be likely pathogenic. However, Sanger sequencing showed that the mother is a carrier, yet the father does not carry the mutation, therefore complicating genetic counseling on the family and subsequent design of PGD on the proposed IVF.

### Long-read sequencing by Oxford Nanopore technology

Due to the presence of Mendelian inconsistency on the c.326G>A mutation, and that sperm sequencing on the father was inconclusive, we suspected that a SV may have encompassed the exon but with breakpoints in non-exonic regions, evading detection by whole-exome sequencing. Due to the complex genomic architecture around this gene, we decided to sequence the patient by low-coverage long-read sequencing on the Oxford Nanopore sequencing platform. Genomic DNA was extracted and large insert-size libraries were created according to the manufacturer recommended protocols (Oxford Nanopore, UK). Five μg genomic DNA was sheared to ~5-25kb fragments using Megaruptor (Diagenode, B06010002), size selected (10-30kb) with a BluePippin (Sage Science, MA) to ensure removal of small DNA fragments. Subsequently, genomic libraries were prepared using the Ligation sequencing 1D kit SQK-LSK108 (Oxford Nanopore, UK). End-repair and dA-tailing of DNA fragments according to protocol recommendations was performed using the Ultra II End Prep module (NEB, E7546L). At last, the purified dA tailed sample, blunt/TA ligase master mix (#M0367, NEB), tethered 1D adapter mix using SQK-LSK108 were incubated and purified. Libraries were sequenced on R9.4 flowcells using GridION X5. Four GridION flowcells generated 2,251,269 base-called reads containing 35,595,548,336 bases with an average read length of 16,579 bp. We used NGMLR ^34^ to align the long reads to the human reference genome (GRCh37). Structural variations (SVs) were called by Sniffles ^34^ and single nucleotide variants (SNVs) were called by Samtools ^35^ for comparison to WES results. Ribbon ^36^ and IGV ^37^ were used to visualize the alignment results and manually validate possible SV calls.

### Sanger sequencing and quantitative real-time PCR

Sanger sequencing and quantitative real-time PCR (RT-PCR) were conducted to validate the long-read sequencing result in the family. Genomic DNA was extracted from peripheral leukocytes using QIAamp^®^ DNA Blood Mini Kit (Qiagen, Germany). The exons of the G6PC gene as well as the potential breakpoint junction were amplified using PCR primers (Table 1). Each PCR reaction was performed in a total volume of 40 μl containing 20 μl of GoTaq^®^ 2 × Green Master Mix, 0.8 μl of 10mM/l mixture of forward and reverse primers, 1.5 μl of 80 ng/μl genomic DNA template, and 17.6 μl of nuclease-free water. The PCR was conducted under the following cycling conditions: 95 °C for 5 min, 35 cycles of denaturation at 95 °C for 30 s, annealing at 58 °C for 30 s, and elongation at 72 °C for 30 s followed by a final elongation of 5 mins, and PCR products were sequenced using Sanger sequencing. The exons of the G6PC gene were also quantitatively tested using quantitative RT-PCR (Table 1). Data were analyzed using the ΔΔCT method.

We further designed an easy assay to detect the deletion reliably by PCR, and performed nucleic acid gel electrophoresis and used the ß-Globin gene as an internal control to validate this assay. The amplification primers for the ß-globin gene are: Primer ß-F: 5′-TGAGTCTATGGGACGCTTGA-3′ and Primer ß-R: 5′-ATCCAGCCTTATCCCAACC-3′. The primers designed to amplify the sequence near the breakpoint were G6PC-DEL-F: 5′-GAGTTAGAAGGAGATGGCGGG-3′ and G6PC-DEL-R: 5′-GGCCTATCCTACATATTAATAGTT-3′ which generates a target fragment of 418 bp.

### Assessment of pathogenicity of the mutations

We analyzed the WES data through the variant filtering pipelines implemented in the ANNOVAR software ^38^, by focusing on coding variants and by removing common variants observed in the gnomAD database. We identified a novel and homozygous missense mutation (c.326G>A; p.C109Y) in the G6PC gene. The novel missense mutation has not been recorded in the HGMD version 2017.4 (http://www.hgmd.cf.ac.uk/ac/index.php) or in the dbSNP (https://www.ncbi.nlm.nih.gov/snp/) database. According to the ACMG-AMP 2015 Standards and Guidelines ^26^, and facilitated by the InterVar software tool ^25^, we analyzed the pathogenicity of the novel mutation and determined that it is a likely pathogenic mutation responsible for the disease manifestation. No other candidate genes were found in our WES analysis that may explain the observed phenotypes of the proband.

### In vitro fertilization (IVF) and pre-implementation genetic diagnosis (PGD)

The parents of the proband had genetic counseling at the Reproductive & Genetic Hospital of CITIC-Xiangya, and proceeded with in vitro fertilization (IVF). A total of four oocytes were retrieved; all were in metaphase II, and all of them were inseminated (day 0) by intracytoplasmic sperm injection. The embryos were cultured to blastocyst stage, and 2-3 zona pellucida cells were used to detect mutations in the G6PC gene, and the embryos were also scored according to the Istanbul consensus (Table S1). PGD was performed using assays designed for detecting both the missense mutation and the exonic deletion in G6PC. The best embryo was selected for implementation after PGD.

## Acknowledgments

We thank the patient and his family members for participating in this study. We thank the PGD group members and the IVF group members at the Reproductive & Genetic Hospital of CITIC-Xiangya, who worked together to complete the project. This work is supported by the Reproductive & Genetic Hospital of CITIC-Xiangya and Central South University.

